# TLR9 activation in large wound induces tissue repair and hair follicle regeneration via γδT cells

**DOI:** 10.1101/2024.02.15.580480

**Authors:** Xinhui Li, Yang Yang, Zumu Yi, Zhaoyu Xu, Shuaidong Chen, Tiantian An, Feng Zhou, Chen Deng, Yi Man, Chen Hu

## Abstract

The mechanisms underlying tissue repair in response to damage have been one of main subjects of investigation. In this study, we leveraged the wound-induced hair neogenesis (WIHN) models in adult mice to explore the inner correlation. Our investigation revealed that heightened release of mitochondrial DNA (mtDNA) accompanying tissue damage activated the toll-like receptor 9 (TLR9) pathway, influencing the repair process and the ultimate number of regenerated hair follicles. Furthermore, our analysis of single-cell RNA sequencing comparisons demonstrated increased TLR9 activation was associated with the recruitment of gamma delta T cells (γδT). Inhibition of γδT cell recruitment led to a reduction in the population of γδT cells and a more fibrotic healing outcome. Notably, these γδT cells exhibited distinctive high production of AREG, contributing to the rapid increase of local AREG levels around the epidermis and influencing the fate commitment of keratinocytes. These findings provide new insights into the roles of TLRs as critical mediators in the sense of tissue damage, the modulation of immune cell activity, and the ultimate influence on healing outcomes.

**Teaser:** Starting with how tissue injury stimulates downstream tissue repair and regeneration through relevant signals, this study explored the phenomenon and correlation between tissue damage and TLR9, and the effect of TLR9 on γδT, keratinocytes and the healing outcomes.

## Introduction

Scar-free healing and functional regeneration after tissue injury have long been desired goals. Intriguingly, a rare capability for hair follicle (HF) regeneration in adult mice and rabbits has been observed in large full-thickness wound centers in the wound-induced hair follicle neogenesis (WIHN) model(*1, 2*).

Tissue repair and regeneration after injury are closely related to inflammatory processes. The exceptional regenerative capacity of fetal tissue is often attributed to its underdeveloped immune system and the detrimental impact of immune reactions on the process of regeneration(*3*). Nevertheless, in the fully matured immune system, the relationship between inflammation signals and regeneration may exhibit dissimilar characteristics(*4, 5*). Damage associated molecular patterns (DAMPs) serve as signaling molecules that indicate tissue injury, thereby initiating the regenerative process (*6*). However, the precise mechanisms through which these factors influence the immune system, the fate of stem cells, and the neogenesis of hair follicles remain incompletely understood. Interestingly, previous investigations have documented a positive correlation between the size of wounds and their regenerative potential, whereby larger or more severely damaged wounds elicit more robust regeneration(*1, 7*).

In some literature reports, excessive or prolonged inflammatory response can cause fibrosis (*4, 8, 9*); while in others, a stronger inflammatory response may activate a more robust regenerative response(*10, 11*). Therefore, we hope to explore how the inflammatory response will change under enhanced tissue damage, and how this stronger inflammatory response affects immune cell and regenerative behavior.

Our previous research demonstrated that the implementation of an aligned extracellular matrix (ECM) scaffold resulted in improved regeneration of hair follicles, accompanied by an immunomodulatory impact (23, 24). Furthermore, our study delved into the crucial involvement of adaptive immune cells in the process of extensive wound healing and regeneration(25). However, more researches are needed to explore how multiple immune cells are affected by signals derived from tissue damages and then initiate and coordinate tissue healing and especially regeneration related reactions. Here, we began with the healing outcomes changed with various degrees of damage. We found the expression of TLR9 was elevated as the damage worsened. Moreover, we further studied the function and effects of TLR9 on multiple immune cells, especially in γδT cells. The prominent and profound immunoregulatory effects were found after activation of TLR9. The expression and activation of TLR9 in monocytes and macrophages affected the migration of γδT cells, which have been proved to have impacts on regeneration of skin wounds. Unveiling the immune cell composition and function changes affected by TLR9 provided a novel and further understanding of damage induced tissue repair and regeration.

## Results

### Tlr9 is activated by the mtDNA Released by Tissue Damage

We refer to the classic WIHN model, which involves creating a large wound on the back of mice and allowing it to heal naturally without splinting. In order to explore the tissue changes under the relatively strong degree of tissue damage, we enlarged the diameter of the wound to 1.8cm. To eliminate the asynchrony of wound healing caused by different healing speed under different situations, we used the second day after re-epithelialization (SD2) as a detection window phase for hair follicle regeneration, according to relevant literature, hair follicles begin to form at around SD 0-7 days(*12, 13*) (Fig 1A). To investigate the impact of wound severity on regeneration outcomes, we made the enhanced injury on large wound (enhanced_LW) model in study of Nelson et al(*14*) by adding 6-8 short radial incisions around the circular wound, which increased the degree of wound damage without changing the wound area. Since many studies suggest that mechanical tension and other factors may also affect the degree of regeneration, we designed various types of enhanced large wound models to rule out the possibility of these confounding factors (Fig S1A). By comparing various wound types, we found multiple types of wounds could enhance hair follicle regeneration compared to standard large wounds (WT_LW) at 28 days post wounded (PWD28) (Fig S1B).

**Fig. 1.**
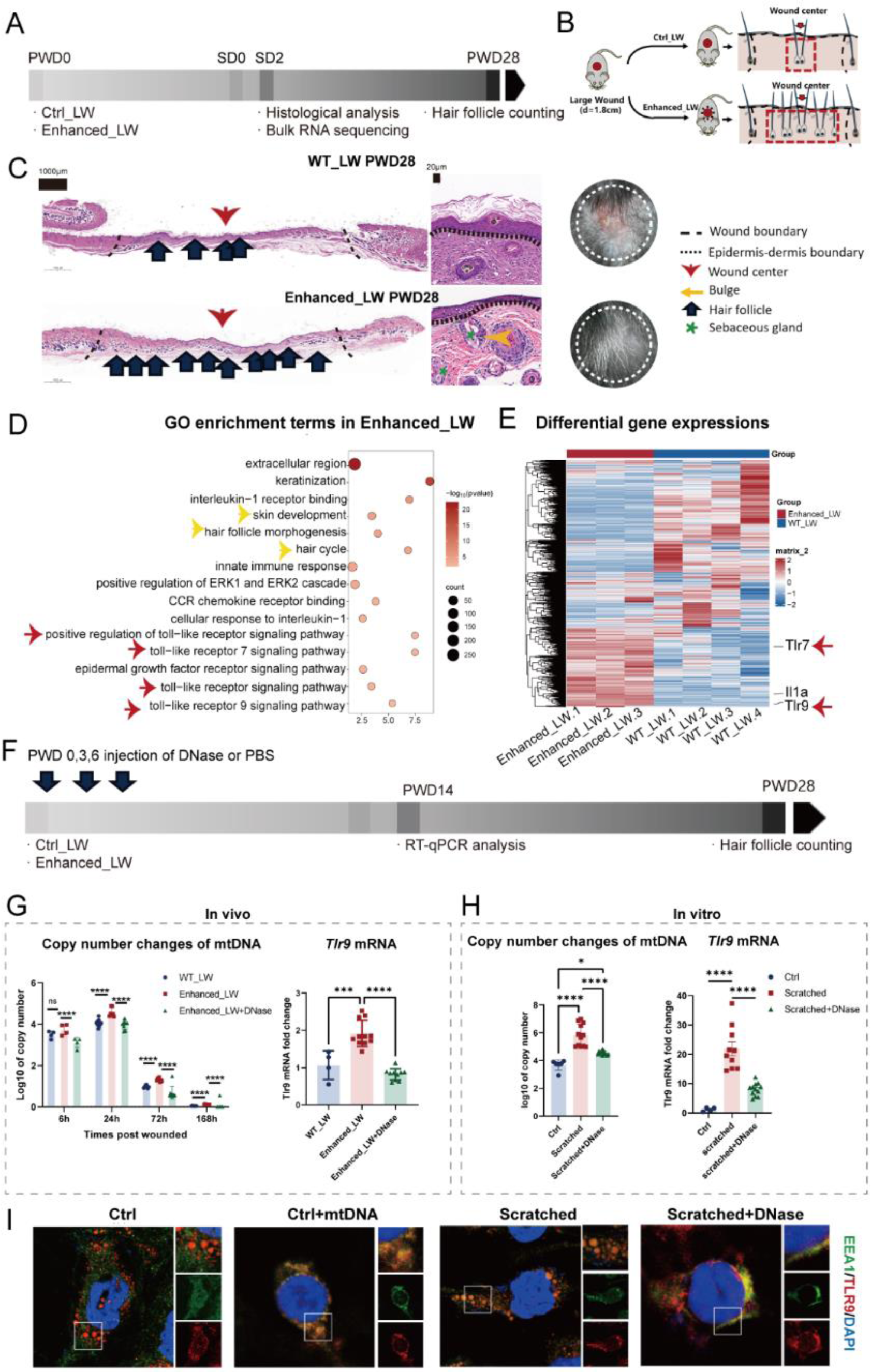
Evaluation of the wound healing process with enhanced_LW. **(A)** Workflow for evaluating large-scale wound healing. **(B)** Surgical processes for skin wound models. **(C)** Representative H&E images and appearances of WT_LW and enhanced_LW at PWD28. **(D)** GO enrichement terms up-regulated in enhanced_LW compared with WT_LW. Yellow arrows are terms related with skin and hair follicle development. Red arrows are terms related with toll-like receptor signaling pathway. **(E)** The differential expressed genes between WT_LW and enhanced_LW. **(F)** Workflow for enhanced_LW injected with DNase. **(G)** The copy numbers of mtDNA (using Cytochrome B as the representative) performed by quantitative PCR. And the qRT-PCR mRNA expression of *Tlr9*. Statistical analysis was performed using one-way ANOVA with Dunnett’s multiple comparisons test. **(H)** The in vitro tests of copy number changes of mtDNA and *Tlr9* mRNA fold changes at 24h after being scratched. n=3. Statistical analysis was performed using one-way ANOVA with Dunnett’s multiple comparisons test. **(I)** The immunofluorescent staining of TLR9(red) and EEA1(marks endosomes) (green) for control M0-THP1 (treated with PMA for 24h) and M0-THP1 treated with 100ng/ml mtDNA; M0-THP1 treated with 4-5 scratches; M0-THP1 treated with 4-5 scratches and 1μg/ml DNase. All error bars ±SD. *p < 0.05, **p < 0.01, ***p < 0.001, and ****p < 0.0001.

To further explore the relationship between injury severity and hair follicles, and to dissect the biological effects of enhanced injury on the skin, we collected the tissues at wound center of enhanced_LW and WT_LW at SD2 and conducted bulk-RNA sequencing (bulk-RNA seq) (Fig 1A, Fig S1C). The gene ontology (GO) enrichment analysis showed that multiple TLRs family pathways were upregulated in enhanced_LW compared with WT_LW, especially TLR7 and TLR9, as observed by qRT-PCR(Fig 1D-E, Fig S1D-G). TLRs are critical components of PRRs in mammals, capable of recognizing both damage-associated molecular patterns (DAMPs) and pathogen-associated molecule patterns (PAMPs)(*15*). While TLR7 has been extensively studied for its effects on skin due to its use as a modeling agent for psoriasis(*16*), little is known about the relationship between TLR9 and skin wounds. Some studies suggest that TLR9 activation in plasmacytoid dendritic cells (pDC) following tape stripping-induced skin damage can have positive effects on wound healing(*17*). However, whether TLR9 activation may affect hair follicle regeneration or fibrosis and the detailed mechanisms remains unclear. Therefore, we aim to investigate TLR9 in greater depth. TLR9 represents the first receptor discovered to recognize unmethylated CpG DNA, typical of bacterial DNA and mtDNA(*18*), which brought us to mtDNA. MtDNA, being structurally analogous to bacterial DNA, is characterized by the presence of numerous unmethylated DNA regions referred to as CpG islands. Mitochondrial DNA is generally known as an immune stimulatory molecule and is released from necrotic cells after traumatic injury or surgery to activate TLR9, stimulator of interferon genes (STING) and so forth (*19*).

Therefore, we firstly detected the copy numbers of mtDNA in two groups and found that the copy numbers of Cytochrome B (CytB) in mtDNA increased after tissue damage (Fig 1G). Injecting DNase I at the wound bed at PWD 0,3,6 decreased the detected copy numbers of mtDNA, mRNA expression levels of TLR9, and hair follicle regeneration (Fig 1F). To prove that mtDNA affects hair follicle regeneration through TLR9, we extracted mtDNA from mouse liver and injected DNAse I to clear mtDNA after injury on D0, D3, and D6. Hair follicles decreased after clearance, while injecting mtDNA into circulation increased hair follicles, which were inhibited by a TLR9 antagonist (Fig 2D).

**Fig 2.**
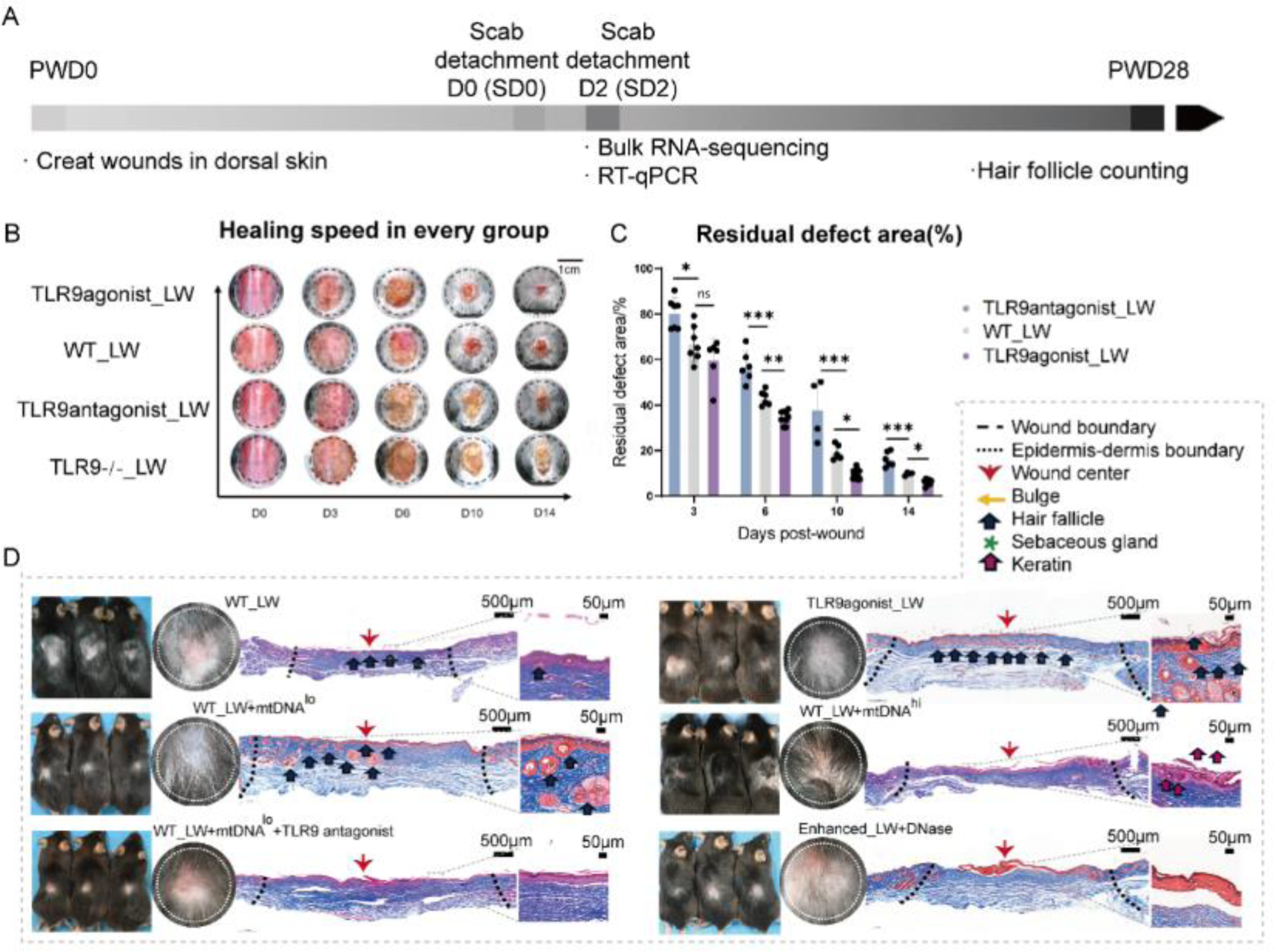
The influence of TLR9 in tissue repair and regeneration. A) Workflow for evaluating the function of TLR9 and mtDNA in large-scale wound healing. The time around scab detachment (SD) was chosen as the key time of hair follicle neogenesis. B) The healing speed and appearance of every group treated with TLR9agonist ODN2395, Invivogene. 4μg per time per mouse at PWD 0,3,6 (TLR9agonist_LW); Ctrl ODN (ODN2395 control, Invivogene. 4μg per time per mouse) (WT_LW); TLR9antagonist (TLR9antagonist_LW) (ODN2088, Invivogene. 4μg per time per mouse); or wound in TLR9 -/- mice treated with PBS (TLR9 -/-_LW). C)The residual defect area percentage for every group. n=4-8 for each group. D) The Masson staining sections and appearance of wounds for every group. The photos and sections were collected at PWD28. All error bars ±SD. *p < 0.05, **p < 0.01, ***p < 0.001, and ****p < 0.0001.

To more intuitively study the association between cellular damage, mtDNA, and TLR9, we simulated tissue damage through scratch experiments using macrophage cells cultured in vitro. We found that the copy numbers of cytochrome b (CytB) and the expression levels of TLR9 increased after 12 hours of cell scratch (Fig 1G). TLR9 is usually located in the endoplasmic reticulum of the cytoplasm and then transferred to endolysosomes after stimulation(*20*). Therefore, we performed IF staining on cells cultured in vitro and found that TLR9 was co-localized with endosomal marker EEA1 after scratch-induced cellular damage. This process requires mtDNA participation (Fig 1I).

### TLR9 activation influence wound healing and hair follicle regeneration

After elucidating the association between injury and TLR9, we want to investigate whether the enhanced hair follicle regeneration in enhanced_LW was caused by activation of TLR9. To explore the effects of TLR9 on wound healing and HF regeneration, we simulated the early release of mtDNA after injury and injected the TLR9 agonist ODN 2395 (4μg per injection, InvivoGen, America) or the TLR9 antagonist ODN 2088 (4μg per injection, InvivoGen, America) at the wound bed on PWD0, 3, and 6. The workflow for evaluating large wound healing is summarized in Figure 2A. The TLR9 agonist significantly accelerated the healing speed (Fig 2B, C). At PWD28, increased HF regeneration in the central area of the wound was observed through tissue alkaline phosphatase (ALP) staining and hematoxylin-eosin staining (H&E staining) (Fig 2D). The regenerated hair follicles exhibited similar morphology and density to the surrounding normal skin, with mature hair bulge and sebaceous gland. In contrast, the wound in TLR9-/-mice or injected with TLR9 antagonist slowed down the healing speed and totally eliminated the HF regeneration (Fig 2D). Therefore, we inferred that TLR9 might play an essential role in wound healing and HF regeneration in large wounds.

### The immune changes caused by TLR9 activation on large wound healing and HF regeneration was investigated by single-cell RNA sequencing

In order to investigate the specific characteristics influenced by the Toll-like receptor 9 (TLR9) agonist during the process of wound healing, we employed a technique known as single-cell RNA sequencing (scRNA-seq) to compare the expression profiles of genes between two distinct groups. Tissue samples were collected from the center of the wound (φ=5mm) on the second day after injury, comprising two groups: WT_LW and TLR9agonis_LW, each consisting of six mice.

These samples were subsequently subjected to analysis using the 10x scRNA-seq platform (Fig 3A). Following cell filtering, unsupervised clustering utilizing Seurat software assigned cells into distinct clusters based on their global gene expression patterns. This was followed by the categorization of these clusters into primary cell classes at the first level. Ten cell types were defined: T cells (Tc), fibroblasts (Fib), myeloid cells (Myl), keratinocytes (Ker), pericyte cells (Perc), Neural crest-derived cells (Neur), endothelial cells (Endo), other cells (Others), neutrophils (Neu), lymphatic endothelial cell (Lyen) (Fig 3B). The composition of each main cluster was listed so that the proportion of cells from two groups could be identified across all cell clusters. Marker genes for each main cluster were shown in the heatmap and listed in Fig 3C. Subsequently, we performed enrichment analysis of overall gene expression in both groups. GO functional enrichment revealed significant up-regulation of many entries related to hair follicle development/epithelial tube morphogenesis/gland and appendage morphogenesis in the TLR9agonist_LW group, as well as hair follicle stem cell-related genes such as Krt17(*21*)、 Sox9(*22*)、Msx2(*23*)、Lgr5(*24*)(Fig 3D,E). In contrast, the WT_LW group mainly up-regulated genes associated with myeloid cell differentiation, phagocytosis, WNT and TGFβrelated terms (Fig 3D,F), which suggested that hair follicle regeneration in the TLR9_agonist group might not be WNT-dependent.

**Fig 3.**
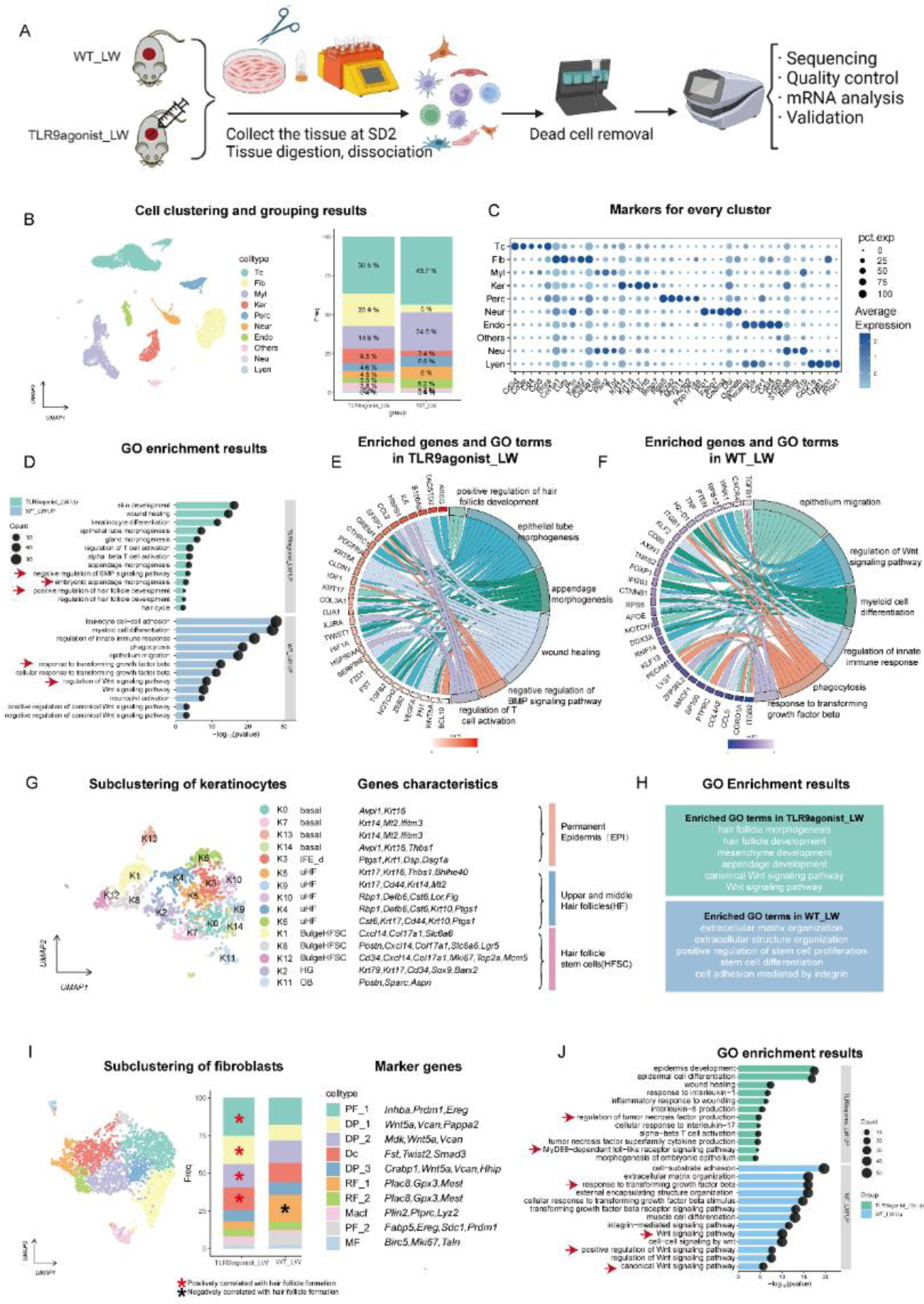
The single-cell atlas of the TLR9 activated microenvironment. A) Schematic for generating scRNA-seq data from large area excisional wounds on SD2. B) Clustering of all cells, showing ten subsets from the two samples. C) The marker genes for each subset are listed. D-F) The enriched terms results and relative genes in each term. G) Subclustering of keratinocytes and the annotation, and ratios and marker genes of every subset. H) The GO enrichment terms up-regulated in TLR9agonist_LW and WT_LW are showed respectively. I) The subclustering of fibroblasts and maker genes. The Red asterisk marks subsets reported positive with the regeneration of skin wound and hair follicle. The black asterisk marks subsets reported positive with the formation of fibrosis or fibrotic heal. J) The GO enrichment terms upregulated in two groups of fibroblasts are showed respectively.

To better analyze the mechanisms underlying the effects of TLR9 activation on wound healing and HF regeneration, we first annotated the two most important stromal cell populations, keratinocytes and fibroblasts, based on previously reported markers. Keratinocytes were divided into permanent epidermis (EPI), upper and middle hair follicles and cells defined as or highly related with hair follicle stem cells (HFSC) (Fig 3G). The subclusters of keratinocytes were further divided and annotated according to markers reported before(*25–28*). The GO enrichment analysis also showed genes related with hair follicle morphogenesis/hair cycle and skin development were upregulated in TLR9agonist_LW, while genes related with extracellular matrix organization and stem cell differentiation enriched in WT_LW (Fig 3H). Furthermore, in order to clarify the effects on keratinocytes and the resulting changes more clearly, we conducted detailed analysis of the development trajectory of keratinocytes and their interaction with immune cells, which can be seen in Figure 5 and 6.

Similarly, we specifically focused on the main clusters that were identified as fibroblasts and subjected them to a secondary round of clustering. Fibroblasts play a crucial role as the primary mesenchymal cells in the dermal layer of the skin, and different subclusters of fibroblasts are spatially distinct with significant functional diversity. Generally, dermal fibroblasts originate from several distinct lineages: (1) the upper lineage consists of papillary fibroblasts (PF), which are in direct contact with the epidermis and contribute to the dermal component of hair follicles; (2) the lower lineage comprises reticular fibroblasts (RF), responsible for synthesizing most of the extracellular matrix (ECM) proteins, and lipo-fibroblasts (LF), which give rise to preadipocyte progenitors in the hypodermis.

Additionally, certain populations of fibroblasts play unique roles in WIHN and are therefore of particular interest to us, namely dermal papilla (DP) and dermal condensate (Dc), which possess unique transcriptional characteristics along with the general fibroblast population (*29*). DP is located at the base of mature hair follicles and serves as the principal signaling niche regulating hair follicle activities(*30, 31*). Dermal condensate (Dc), originating from the papillary fibroblasts, is believed to be the progenitor of DP during embryonic development. In our dataset, we defined six types of fibroblasts consisting of ten subclusters based on previously established marker genes: Inhba+Prdm1+Ereg+ Fabp5+Sdc1+ representing papillary fibroblasts (PF), Pappa2+Mdk+Wnt5a+ Hhip+Vcan+ indicating dermal papilla (DP), Fst+Twist2+Smad3+ characterized dermal condensate (Dc), Plac8+Gpx3+Mest+ indicating reticular fibroblasts (RF), Plin2+Ptprc+Lyz2+ indicating myeloid-derived adipocyte progenitors (Macf), and Birc5+Mki67+Tagln+ representing myofibroblasts (MF) (Fig 3I).

In the WT_LW group, the initial phase of dermal repair was mediated by the lower lineage fibroblasts, particularly the reticular fibroblasts, which were associated with the organization of extracellular matrix organization, external encapsulating structure organization, as revealed by gene ontology enrichment analysis (Fig 3J). In addition, the fibroblasts in WT_LW highly enriched terms of transforming growth factor beta (TGF-β) receptor signaling pathway, which was known as associated with fibrosis(*32*). In contrast, the TLR9agonist_LW wounds exhibited a higher proportion of upper lineage Crabp1+Prdm1+ papillary fibroblasts, which are known to support hair follicle initiation. Interestingly, we also identified the presence of Fst+ Twist2+Smad3+ dermal condensate (Dc) cells in this dataset. During embryonic hair follicle development, Dc acts as a signaling niche that promotes epithelial placode growth and subsequently contributes to hair follicle morphogenesis. In GO enrichment analysis, we observed the genes related with morphogenesis of embryonic epithelium, regulation of tumor necrosis factor (TNF) production were highly regulated in TLR9agonist_LW, which reminded us of the positive correlation between TNF-α(*33*) or IL-1β(*12*) with hair growth in wounds. Additionally, as observed by Gay et al(*9*)., the WNT signaling pathway was also more significant in low-regeneration group at similar late wound healing, after the end of re-epithelialization (Fig 3J).

Given the significantly increased proportion of specific cell populations and enrichment terms in the TLR9agonist_LW group during the critical period of hair follicle regeneration, it is plausible to suggest that they may provide an adequate mesenchymal component for subsequent hair follicle formation in the TLR9-activated group.

TLRs are expressed in antigen-presenting cells, establishing a crucial link between pathogen recognition and the activation of both innate immune effector mechanisms that restrict pathogen replication and adaptive immunity initiation(*34*). To comprehensively understand the impact of TLRs on the host’s overall immune response, we conducted an analysis of various immune cell types using single-cell RNA sequencing.

Firstly, neutrophils, as the primary responders and critical mediators of the innate immune system, play a vital role in the recruitment and differentiation of monocytes(*35*), particularly during the early stages of immune response. In our scRNA sequencing data obtained from the late stage of wound healing, we observed that most subclusters of neutrophils exhibited comparable proportions in both experimental groups (Fig S3B). Macrophage-monocytes and dendritic cells are the main cells expressing TLRs and equipped to coordinate the activation of other immune cells as well as tissue repair(*17*). Five types of macrophage-dendritic cells were identified, according to markers reported in literature(*36*): Inhba+Ptgs2+ Mmp12+anti-inflammatory macrophages (PIM), Ccl8+Mrc1+ Folr2+Fcgr1+ pro-inflammatory macrophages (AIM), Irf7+Il3ra+ plasmacytoid dendritic cells (pDC), Cd74+Cd86+Cd207+Rgs1+ monocyte-derived dendritic cell1 (mDC), Cd8a + Cadm1+ Cadm3+ Clec10a+ Cpvl + conventional dendritic cell1 (cDC)(Fig S3D).As speculated, TLR9 agonist, as a type of pro-inflammatory agent, increased the presence of PIM in wounds while decreasing AIM (Fig S3C). Functional enrichment analysis of the two groups revealed that the TLR9 agonist upregulated numerous items related to protein synthesis and secretion in the large wound group, In consistent with the macrophage activation response to TLRs as reported in (*34*). Further subcluster comparisons and enrichment analysis of macrophage-dendritic cells demonstrated that macrophage-dendritic cells were stimulated and activated in the TLR9_agonist group. pDC, as one of the cell types expressing TLR9 and recognizing self-nucleic acids, have been shown to quickly migrate into wound and produce proinflammatory cytokines such as type I interferon (IFN). However, we did not observe an increase of number of pDC in the late-stage wound of TLR9agonist_LW. We suppose this attributed to rapid recovery of pDC number(*37*). However, in the TLR agonist group, pDC showed significant upregulation of pathways such as toll-like receptor signaling pathway and cytosolic DNA-sensing pathway, as well as activation-related pathways like NF-kb and MAPK (Fig S3E).

WNT pathway, as one of the most important pathways involved in growth, development, and hair follicle regeneration, is also closely related to macrophages (38). Macrophages are known to have the ability to secrete WNT ligand WNT3a and involve in WNT pathway signaling after tissue damage, which has been found correlations with both tissue regeneration and fibrosis(*38*). As a result, we hypothesized that TLR9 promotes hair follicle regeneration by stimulating the production of WNT ligands in macrophages. However, contrary to our initial assumption, macrophages (Fig S3F), and fibroblasts (Fig 3J) in the TLR9agonist_LW exhibited decreased mRNA levels of WNT ligands including *Wnt2, Axin2, Lef1, Tcf3,* and *Ctnnb1* while showing elevated mRNA levels of WNT pathway antagonists like *Sfrp2, Sfrp4.* This discovery reminded us of Gay et al.’s study (9), which highlighted the detrimental effects of macrophage over-phagocytosis, resulting in the engulfment of the WNT antagonist Sfrp4. The excessive activation of the WNT pathway ultimately inhibited hair follicle development (9). Moreover, our functional analysis of macrophage phagocytosis revealed lower expression levels of genes associated with phagocytic function in the TLR9agonist_LW compared to the control group (Fig S3G).

Expression of TLRs in antigen-presenting cells can also lead to the initiation of adaptive immunity. In our single-cell data, T cells were found to be the most abundant adaptive immune cells (Fig 3B). Our previous research has shown that T cells play a crucial role in wound healing and hair follicle regeneration(*9*), and rag-/-mice lacking mature T and B cells lose the ability to regenerate hair follicles in the WIHN model(*39*). Therefore, we further investigate the role of T cells and how they are influenced by tissue damage or TLR9, and how they ultimately affect wound healing and hair follicle regeneration outcomes. Subclustering of T cells resulted in five main subsets including CD4-Il17a+ Cd7+ γδT cell (γδT), CD4+Ifng+Il18+ T helper 1 (Th1), CD4+Gata3+Il4+ T helper 2 (Th2), Foxp3+Ctla4 + T regulatory cell (Treg), Cd8b1+Nkg7+ cytotoxic T cell (CTL), based on markers from published research(*40*)(Fig 4A). After defining cell subsets, we observed a noteworthy increase in the number of γδT cells and numerous enriched GO terms related with activation and differentiation of γδT cells in the TLR9 agonist_LW group (Fig 4A, C). The increased γδT cells were primarily IL17-producing γδT cells from the dermis, characteristic of Cd27-Cd44+ Il17a+ Il7r+ Blk+ Maf+ Rorc+(*41*). The increase in γδT cells was also confirmed by flow cytometry (Fig 4D, E), indicating that TLR9 stimulation may significantly affect γδT cells. Additionally, similar to TLR9 agonists, enhanced tissue damage also resulted in an increase in the number of γδT cells, further suggesting that γδT cells are influenced by the degree of tissue damage and the activation of TLR9. To explore the source of the increased γδTcell numbers, we scored the proliferation-related genes of γδTcells with AddModuleScore in Seurat and found that the cell proliferation genes of γδT cells in the TLR9 stimulated group didn’t increase, indicating the increase of γδTcells in wound might caused by enhancement of migration rather than proliferation (Fig 3F).

**Fig 4.**
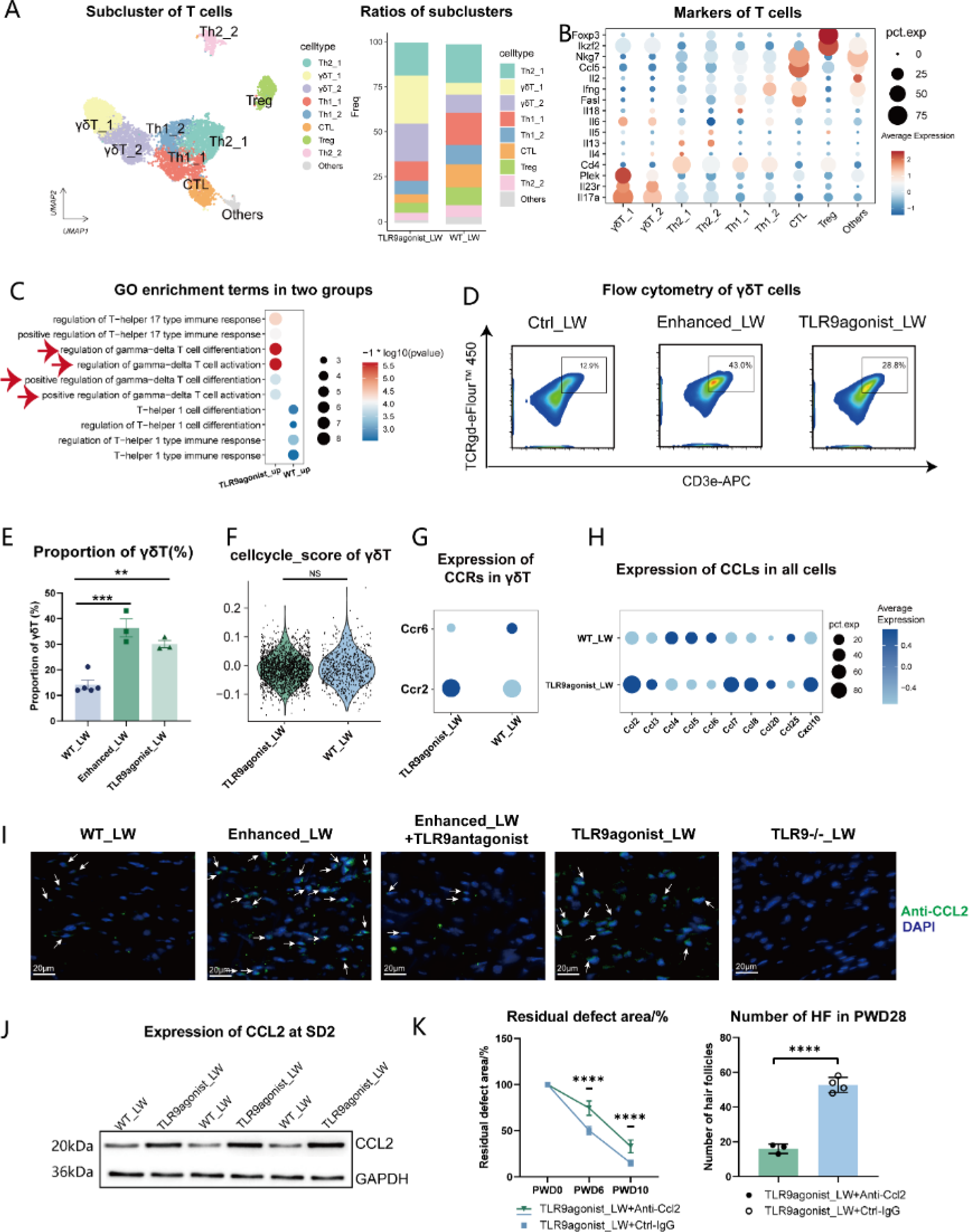
The changes in cell proportion and function of T cells in TLR9 activated skin wounds. A) The subcluster of T cells and the proportion of each subset in the two groups. B) Markers used for identification of cell types. C) The GO enrichment terms in two groups. D-E) The flow cytometry results of γδT cells in three groups. n=3-5 for each group. F) The gene expression scores for cell cycles in γδT cells of two groups. The list of genes can be seen in Fig.S4 and the protocols of scoring can be found in experimental sections. G) The expression of CCRs in γδT cells of two groups. H) The expression of gene members in CCL family in all cells of two groups. I) Immunofluorescence staining was performed to visualize the expression of the CCL2 protein in the dermal part of skin wounds with different treatments at SD2. Scale bar: 20 μm. J) The expression of CCL2 in wounds at SD2 was detected by Western blotting. GAPDH was used as loading control. K) The healing speed and regenerated hair follicle numbers in wounds treated with anti-CCL2 antibody (26161-1-AP, Proteintech)(10 μg per time for per mouse) at PWD 0,3,6 or control-IgG from same species. n=4 for each group. All error bars ±SD. *p < 0.05, **p < 0.01, ***p < 0.001, and ****p < 0.0001.

Due to the constant migration and movement of γδT cells between lymph nodes and peripheral tissues, their migration speed and pattern undergo changes in cases of tissue infection or injury. We therefore investigated whether the increase in γδT cell numbers result from an increase in their migration into the skin. We found that almost all γδT cells were CCR2hi CCR6lo cells (Fig 3 G,H). In steady-state conditions, CCR6 controls γδT 17 trafficking to the dermis; however, in cases of tissue damage, CCR2 controls the rapid migration of γδT 17 to damaged sites(*42*). Through Cellchat analysis of cellular interactions between γδT cells and all other cells, we found among all CCR2 ligands, CCL2 was the strongest acting and expressed one (Fig S3H), significantly upregulated in the TLR9 agonist group. The IF staining and western blot (WB) results also proved that CCL2 was elevated in the TLR9agonist_LW and Enhanced_LW groups (Fig 4I, J). Therefore, we speculated that TLR9 agonist may promote the chemotaxis of γδT cells through CCL2. By injecting anti-CCL2 or Ctrl-IgG at the same time as TLR9 agonist injection, we found the effect of TLR9 agonist on γδT cells required CCL2. At the same time, we also observed that reducing the number of γδT cells at the wound site with anti-CCL2 antagonized the effects of TLR9 agonists on wound healing and hair follicle regeneration (Fig 4K).

### The role of γδT in promoting hair follicle regeneration is achieved through Areg rather than IL17

Although anti-CCL2 reduced the number of γδT17 cells and had a significant inhibitory effect on wound healing and hair follicle regeneration, we were still unsure which molecule was responsible for promoting hair follicle regeneration via γδT cells. It has been previously reported that the promotion of fgf9 secretion by gft17 activates the WNT pathway of keratinocytes(*43*), but the above section had shown that the promotion of hair follicle regeneration by TLR9 agonists was independent of WNT. Further investigation reveals that fgf9 mRNA levels are extremely low in both groups of γδT cells. This suggests that the effect of γδT cells induced by TLR9 agonist stimulation is achieved through a new mechanism. To explore the impact of γδT cells on keratinocytes, we analyzed the cell interactions between T cells and keratinocytes using cellchat (Fig 5A, B). The number of T-ker interactions was significantly increased in the TLR9 agonist group (Fig 5A). Among the significantly increased pathways, IL17 was the most prominent (Fig 5B). Through Cellchat, we identified that IL17 was produced by γδT17 cells and mainly received by bulgeHFSC, IFEB, IFET, and other cells that may be closely related to hair follicle development. By comparing L-R pairs, we found that IL17A was the most influential member of the IL-17 family. Given the reported role of IL-17A in promoting wound healing(*44*) and the stemness of keratinocytes(*45, 46*), we hypothesized that the action of γδT17 cells is also achieved through IL-17A. However, by injecting Anti-IL17A into the wound site early in the experimental period, we found that although IL17A had a significant effect on wound healing speed, antagonizing IL-17A did not affect the hair follicle regeneration promoted by TLR9 agonists(Fig 5G,H). In addition, according to literature, in the WIHN model without intervention, there was no difference in healing outcomes between IL-17a/f -/- mice and WT mice(*12*). Therefore, we speculated that the action of γδT cells is achieved through other signaling molecules.

**Fig 5.**
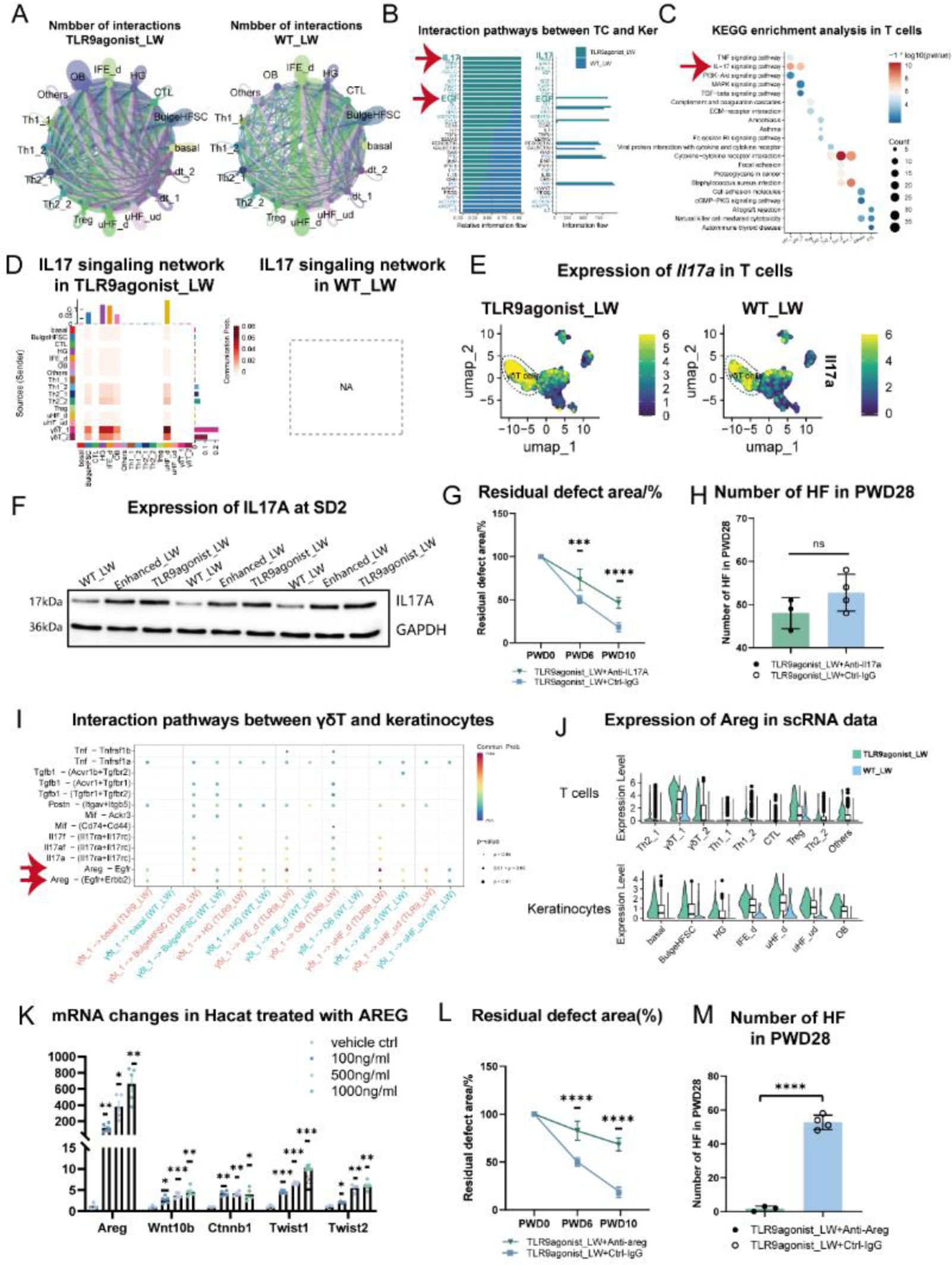
The function of γδT cells are mediated by AREG rather than IL-17A. A) The interaction numbers between T cells and keratinocytes in TLR9agonist_LW and WT_LW. B) The interaction pathways differential in TLR9agonist_LW and WT_LW. C) KEGG analysis showed that the prominent IL-17 signaling pathway in γδT cells. D) The IL-17 interaction networks between T cells and keratinocytes in TLR9agonist_LW and WT_LW. E) The feature plots showing the expression of Il17a in T cells. The main cells expressing Il17a were γδT cells. F) The expression of IL-17A in WT_LW, Enhanced_LW and TLR9agonist_LW at SD2 was detected by Western blotting. GAPDH was used as loading control. G-H) The changes of healing speeds and regeneration outcomes after injection of anti-IL17A (10μg per time per mouse. PAB30184, Bioswamp) or Control-IgG (30000-0-AP, Proteintech) into the wound bed at PWD 0,3,6. I) The bubble diagram showing the pathways mediating the interaction between γδT cells and keratinocytes in two groups. J) The expression level of Areg in γδT cells and keratinocytes respectively. K) The mRNA changes of *Areg, Wnt10b, Ctnnb1, Twist1, Twist2* in Hacat cell after culturing with rh-AREG for 48h. L-M) The changes of healing speeds and regeneration outcomes after injection of anti-AREG (15μg per time per mouse. HY-P77868, MedChemExpress) or Control-IgG (30000-0-AP, Proteintech) into the wound bed at PWD 0,3,6.

The EGF family was also a highly represented signaling molecule in T cell and keratinocyte interactions, and one of the significantly different molecules between the two groups. Bulge HFSC was highly co-localized with EGFR during embryonic development(*47*), and EGFR KO mice exhibit impaired hair follicle differentiation and multiple hair shaft abnormalities, indicating the crucial role of EGF signaling in hair follicle development. Through cellchat L-R scoring, we found that areg was the strongest acting signal in the EGF family. The areg signal was involved in almost all γδT17-related actions on various keratinocytes, with IFEB and bulgeHFSC being the most significant cell populations involved in hair follicle development. By injecting anti-areg monoclonal antibodies to antagonize the effect of areg, we found that wound healing and hair follicle regeneration were significantly reduced, indicating the role of areg(Fig 5L,M). In addition, in order to elucidate the mechanism by which areg acts on keratinocytes, we conducted an in vitro study using human Hacat cells. Various concentrations of recombinant AREG (rh-AREG) were added to the cells, and through RT-qPCR analysis, we observed significant upregulation of genes associated with hair follicle regeneration, such as Wnt10b, Ctnnb1, Twist1, and Twist2.

Concurrently, we also observed a significant increase in the expression level of areg itself upon addition of recombinant AREG, suggesting the presence of positive feedback loop, which was consistent with the results in cellchat where the keratinocytes are recognized as key recipients and responders to AREG. Therefore, we hypothesize that AREG produced and secreted by the γδT cells may act on keratinocytes via paracrine signaling, boosting the production of AREG by keratinocytes themselves, and led to a localized rapid elevation of AREG levels and enhancement of its effects.

### AREG may be associated with the fate commitment of keratinocytes

To elucidate the changes in behavior and differentiation tendency of keratinocytes under TLR9 agonist stimulation, we analyzed the development and differentiation trajectory of keratinocytes in two groups using monocle2 and RNA velocity. Unsupervised clustering of RNA velocity in keratinocytes revealed four major differentiation pathways. Based on cell annotation and spatial localization of annotated cells, we determined four developmental trajectories, namely as: self-renewal of bulgeHFSCs (trajectory 1), regeneration of new hair germs (HG) (trajectory 2), development of hair follicles (trajectory 3), and differentiation of the permanent epidermis (trajectory 4) (Fig 6A). Trajectory 1originated from the highly proliferative Cluster 12 cells characterized by Mki67+ Ccnb1+ Pcna+ and terminated at bulgeHFSC, suggesting that pathway one may represent the proliferation and self-renewal of bulge HFSCs (Fig 6D). Trajectory 2 was initiated by IFEB and culminated in HG characterized by Krt79+Krt17+Sox9+(Fig 6E), which was mainly derived from the development of IFE with lineage plasticity and migration after injury in WIHN (50,51), indicating that trajectory 2 represented the process of generating new HGs in WIHN. Trajectory 3 originated from moderately differentiated keratinocytes and gradually underwent hair follicle differentiation, indicating that trajectory 3 represented the process of hair follicle development (Fig 6I). Trajectory 4 originated from basal-like keratinocytes and underwent gradual differentiation, indicating that trajectory 4 represented the process of differentiation of the interfollicular epidermis.

**Fig 6.**
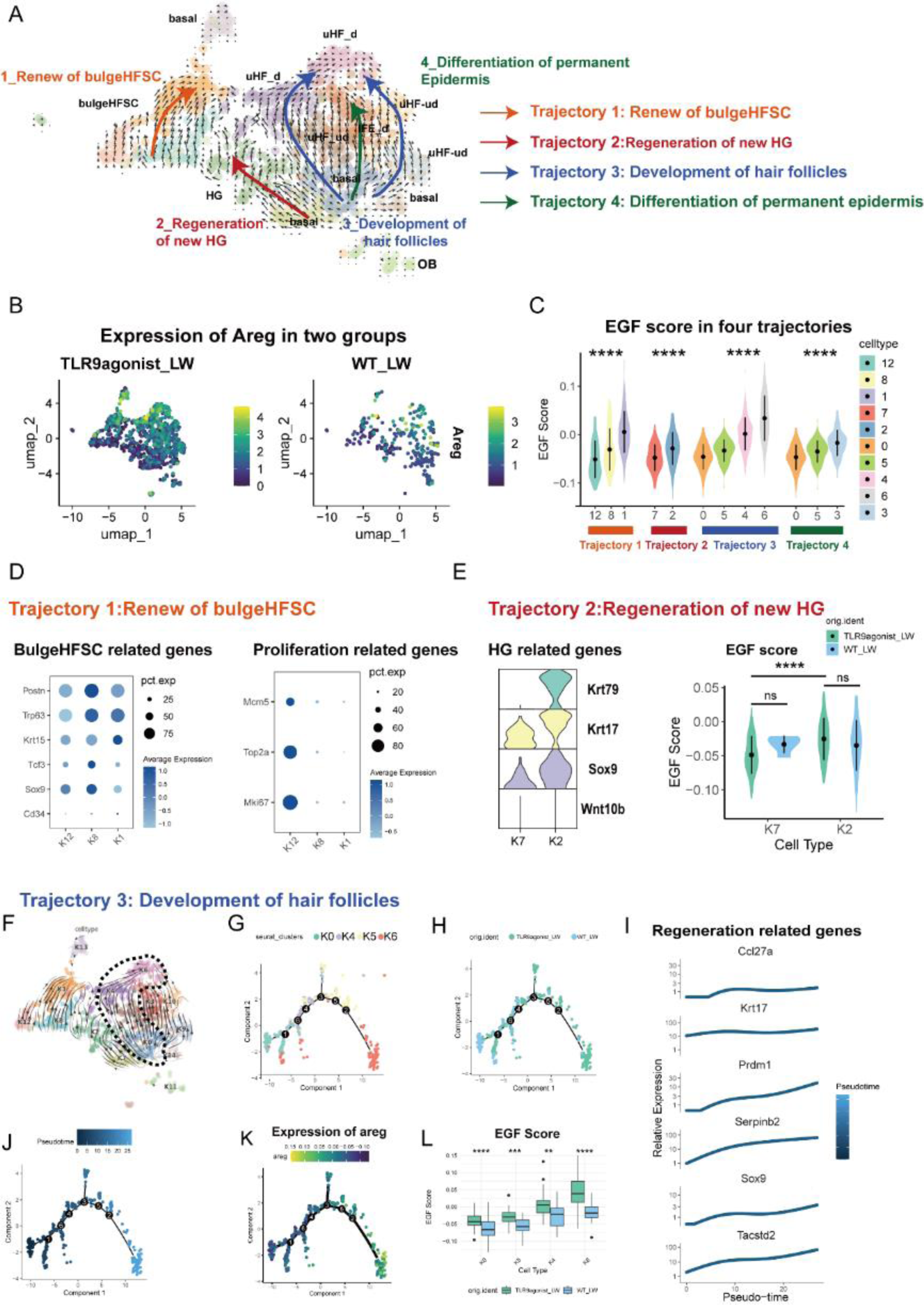
The development trajectories of keratinocytes and the involvement of AREG. A) The RNA velocity analysis of keratinocytes. Trajectory 1: self-renewal of bulgeHFSCs; trajectory 2: regeneration of new hair germs (HG); trajectory 3: development of hair follicles; trajectory 4: differentiation of the permanent epidermis. B) The featureplots of expression of Areg in two groups. C)The scores of EGF in every trajectory. D) The marker genes of bulgeHFSC and proliferation related genes. E) The HG related genes and EGF scores changes in trajectory 2. F-I) The development of hair follicles. G) The pseudotime analysis plotting the sequential changes and transformation between four subclusters in trajectory 3. H) The pseudotime analysis plotting the differential position of cells in two groups. I)The pseudotime analysis plotting expression changes of hair follicle formation and regeneration related genes. J-K) The pseudotime plots and changes of expression levels of areg. L)The EGF scores in trajectory 3. Vlnplot and box plot graphs indicated the value of minimum, first, quartile, median, third quartile, and maximum. Statistics as in Fig. 1.

To elucidate the distinct cellular states during epidermal differentiation and the generation of new hair follicles in the context of WIHN, we initially extracted the major cell clusters (clusters 0, 4, 5, 6) associated with trajectory 3 and investigated the differentiation state of keratinocytes through pseudotime analysis (Fig 6B). As pseudotime progressed (Fig. 6C-E), the expression of regeneration-related markers such as Sox9 and krt17 increased, indicating the developmental pattern (Fig 6E). Concurrently, there was a higher concentration of TLR9_agonist cells in the late stages of the pathway, suggesting a stronger tendency towards hair follicle differentiation (Fig 6D). To understand the role of AREG in hair follicle development, we also performed EGF scoring. During the process of hair follicle development, both the expression of AREG and EGF scoring increased as the developmental pathway advanced, revealing differences between the two groups (Fig 6G, H). These results suggest that increased expression of Areg can maintain the stemness of keratinocytes and promote hair follicle development.

## Discussion

Since the discovery of the WIHN phenomenon, one of the most interesting and urgent phenomena to study is the requirement of sufficiently large wound areas for hair follicle regeneration. These hypotheses encompass a range of factors, including the unique mechanical topology of large wounds, the extent of tissue damage within such wounds, and the varying concentration and spatial distribution of signaling molecules that either promote or hinder regeneration in large wounds. These hypotheses have received partial support through empirical evidence. However, most existing models for studying this phenomenon are susceptible to confounding factors. To address this limitation, our study employed diverse mouse models that encompassed different levels of injury on the dorsal skin. By doing so, we effectively minimized the confounding effects arising from potential synergistic factors. Through this meticulous approach, we conclusively demonstrated the correlation between the degree of tissue damage and the subsequent regenerative outcomes, and unraveled the underlying mechanisms involved.

The recognition of damage-associated molecular patterns (DAMPs) and pathogen-associated molecular patterns (PAMPs) by pattern recognition receptors (PRRs) represents a fundamental mechanism in the initial response to tissue damage or pathogen invasion. In the aftermath of tissue injury, beyond the activation of DAMPs and PAMPs, two pivotal steps must be orchestrated to facilitate tissue repair and regeneration: (1) the coordinated mobilization and activation of pertinent precursor cells, facilitating the reconstitution of damaged tissue structures, and (2) the induction of morphogenetic and regenerative pathways within both precursor cells and stromal cells(*48*). The elucidation of the impact of PPR activation on the immune microenvironment may shed light on this inquiry. Activation of PPRs elicits intricate inflammatory responses, characterized by the recruitment, proliferation, and activation of diverse hematopoietic and non-hematopoietic cells, thereby initiating and coordinating tissue repair and defense responses (*38*).

In this study, we investigated the influence of tissue damage-related signals on the immune microenvironment and the repair process by assessing the number of regenerated hair follicles associated with varying degrees of injury. By analyzing bulk RNA sequencing data from large wounds with different levels of damage, we observed variations in the activation levels of TLRs and specifically TLR9. Previous research has demonstrated the involvement of TLRs, a significant subtype of PPRs, in damage sensing and the initiation of tissue repair and regeneration(*15*). TLRs have also been implicated in the regeneration of skin wounds. For instance, TLR3 activation triggered by the detection of released dsRNA from injured cells upregulates IL-6 expression and induces STAT3 phosphorylation in epidermal keratinocytes(*7, 10*). However, TLR9 has often been overlooked due to its lower expression levels, despite being presumed to be expressed solely in response to skin injury or external stimuli in human and mouse skin. Nevertheless, research on the role of TLR9 in skin wound healing is largely insufficient. In this study, we experimentally confirmed that the substantial increase in hair follicle regeneration caused by enhanced tissue damage largely relies on TLR9. Mice lacking TLR9 or those treated with a TLR9 antagonist shortly after injury failed to regenerate their hair follicles.

Furthermore, we discovered that the enhancement of large wounds compared to ordinary large wounds is primarily facilitated by the release of free mtDNA, which activates TLR9. During cellular trauma, mitochondrial DAMPs, including mtDNA containing CpG DNA repeats, have been observed to be released and activate TLR9, GMPAMP synthase-stimulator of interferon genes (cGAS-STING), and neutrophil extracellular traps (NETs) (*19*). Previous studies have frequently associated mtDNA with NETs, which was demonstrated to impede WIHN, thus implying a negative effect of mtDNA on hair follicle regeneration. Through meticulous in vitro and in vivo investigations, we discovered that lower levels of mtDNA can stimulate TLR9 activation and hair follicle regeneration (Fig 2G). Although the WIHN phenomenon is absent in human skin, mtDNA has also been detected in cases of human tissue damage, accompanied by increased TLR9 expression, demonstrating a similar mechanism. These findings offer novel insights into the activation of signaling receptors implicated in tissue damage, the orchestration of immune responses, and ultimately, the outcome of tissue repair and regeneration. Therefore, we propose the hypothesis that greater tissue damage induces heightened activation of the TLR9 pathway through mtDNA release, which may contribute to WIHN.

Combining single-cell analysis with in vitro and in vivo validation, we observed that TLR9 activation exerts broad and profound effects on various immune cells, particularly facilitating the recruitment of γδT cells, known to play a crucial role in hair follicle regeneration. As one of the key barrier cells residing in the skin, gamma delta T cells (γδT) have been found to play a vital role in detecting skin integrity, maintaining skin homeostasis, aiding in wound repair, preventing infection, and preventing malignant tumors (*49, 50*). Specifically, wounding has been shown to up-regulate epidermal-derived IL-1α, which serves as a potent activator of γδT cells.

Subsequently, activated γδT cells promote hair follicle stem cell (HFSC) proliferation and migration, thereby facilitating injury-induced hair regeneration (*43*) (*51*) (*52*). In addition, γδT cells have also been suggested to be associated with Toll-like receptors (TLRs). The number of γδT cells decreases in skin wounds of TLR3-/-mice (*7*). However, more detailed evidence is needed to elucidate the relevance between the activation of TLRs and T cells, especially γδT cells.

Through analysis of cellular interactions and in vivo and in vitro experiments, we discovered that γδT cells produce AREG, a member of the EGF family, after TLR9 stimulation. This subsequently promotes AREG production by keratinocytes through paracrine signaling, leading to a positive feedback loop that rapidly and significantly increases AREG levels. Our single-cell analysis and in vivo and in vitro research also revealed that AREG affects the expression of genes such as *Twist1, Twist2, Wnt10b*, and *Ctnnb1* in keratinocytes and significantly influences the process of hair follicle development. Although γδT cells have been found to promote hair follicle regeneration by producing FGF9, our samples showed that FGF9 was barely detected in both groups, suggesting that γδT cells may affect regeneration outcomes through other pathways. Additionally, our findings demonstrate that although most of the recruited γδT cells were IL17-producing γδT cells, the addition or antagonism of IL17A did not alter hair follicle regeneration outcomes. Our study not only sheds new light on the role of γδT cells in wound-induced hair neogenesis (WIHN), but also provides explanations for how these cells interact with injury-related signals and ultimately affect the behavior of matrix cells involved in repair.

Our work highlights important future directions for the field. One critical question is whether similar mechanisms exist in humans. As previously discussed, TLR9 expression is up-regulated in human skin after injury, and γδT cells are also present in the human dermis. Moreover, AREG-producing γδT cells have been found in tumor cells. Therefore, it remains to be determined whether γδT cells in human skin will be similarly affected by TLR9 stimulation and produce AREG in response to skin injury. Additionally, aside from dermal γδT cells, will skin injury and TLR9 activation also affect epidermal DETCs or macrophages? DETCs, as one of the critical cells that reside in the epidermis and monitor the status of adjacent keratinocytes through dendritic projections, undergo rapid morphological and functional changes in response to epithelial damage(*53*). Although we were unable to collect sufficient DETCs for analysis due to the limit of single-cell transcriptome cell numbers in this study, the extent of tissue damage and how TLR9 affects DETCs remain important issues worthy of further research. In apart, lower WNT signaling in macrophages and fibroblasts and phagocytosis of macrophages were observed in TLR9 activated group within a few days after re-epithelialization, as reported in gay et al(*9*). Could TLR9 function through similar mechanisms that inhibit late excessive and prolonged WNT activation? In the future, we will explore the detailed mechanisms by which TLRs influence other processes of skin repair and regeneration and to investigate whether any of these mechanisms could explain the healing outcome changes and the lack of WIHN in humans.

## Materials and Methods

### Ethical approval

The experimental procedures undertaken in this study were granted ethical approval by the Institution Review Board of West China Hospital of Stomatology (Approval No. WCHSIRB-D-2023-018).

### Excisional wound model and implantation procedures

Male wildtype C57BL/6 J mice (Dossy Experimental Animals Co., Ltd.) and C57BL/6Smoc-Tlr9em1Smoc mice (TLR9-/-mice) (Cat. NO. NM-KO-190168) ((Shanghai Model Organisms Center Inc., Shanghai, China), aged 6-8 weeks and weighing approximately 20 g, were utilized for the research. Genotyping of the Tlr9 locus was performed using the following primers: 5′-GCTCTCCCACTTTCTCTTCCTCTC −3′ and 5′-TGCCGCCCAGTTTGTCAG −3′ for the wild-type allele, and 5′-TGCCGCCCAGTTTGTCAG −3′ and 5′-ACGGGAAAAGGGTGGGTGTG −3′ for the Tlr9 KO allele. The mice were housed under standard conditions including a temperature range of 21–27 °C, a humidity level of 40–70%, and a 12-hour light-dark cycle with ad libitum access to food. The number of animals utilized for each experiment is specified in the figure legends. Circular full-thickness wounds with diameters of 0.6 cm or 1.8 cm were created on the dorsal skin of the mice. The mice were divided into several groups for further study: large wounds (1.8 cm diameter) treated with PBS (WT_LW) or TLR9 agonist in the wound bed (TLR9agonist_LW), and small wounds (0.6 cm diameter) treated with PBS (WT_SW).

Additionally, to investigate the effects of different degrees of damage, three types of large wounds (1.8 cm diameter) with additional incisions were designed and used: dotted enhanced_LW with 6-8 poke points through the epidermis and dermis, parallel enhanced_LW with 6-8 short incisions parallel to the cutting edge, and vertical enhanced_LW with 6-8 radial short incisions perpendicular to the cutting edge. Mice were euthanized at 1–4 weeks after the surgery, and round full-thickness samples with diameters of 10 mm (for small wounds) or 25 mm (for large wounds) were harvested. To investigate the function of CCL2, IL17A, and AREG, anti-CCL2 (26161-1-AP, Proteintech) at a dosage of 10 μg per mouse, or anti-IL17A (PAB30184, Bioswamp) at a dosage of 10 μg per mouse, or anti-AREG (HY-P77868, MedChemExpress) at a dosage of 15 μg per mouse was injected into the wound bed at postoperative day (PWD) 0, 3, and 6. Rabbit IgG served as the control antibody (30000-0-AP, Proteintech) in all cases.

### RNA isolation and real-time PCR

For in vivo mouse samples, isolation of total RNA was conducted after homogenization using the RNeasy Mini Kit (Qiagen, 74106) as appropriate. RNA concentration and purity were determined using the NanoDrop2000c (Thermo Fisher Scientific). Reverse transcription of RNA into cDNA was accomplished with the PrimeScript RT reagent Kit with gDNA Eraser (RR047A, Takara), followed by quantitative RT-PCR (qRT–qPCR) using the TB Green® Premix Ex Taq™ (Tli RNaseH Plus) (RR420A, Takara). Data were normalized to GAPDH expression utilizing the 2^-ΔΔCT^ method. The standard deviation (S.D.) of samples carried out in technical triplicates was represented by error bars. The supplemental materials contain the primers used for RT-qPCR.

### Cell culture and scratch test

Since TLR9 expression is primarily documented in monocytes-macrophages in existing literature, we utilized TRAIL-resistant human acute myeloid leukemia cells (THP-1) for our research. THP-1 cells were obtained from the cell library of the Chinese Academy of Sciences and cultured in RPMI medium supplemented with 10% FBS, 100 U/mL penicillin, and 100 mg/mL streptomycin at 37 °C in a 5% CO2 incubator. Scratch assays were employed on densely adherent M0-THP1 cells to simulate tissue injury, following the method used by Nelson et al(7). To induce differentiation of THP-1 cells into a macrophage-like phenotype, 106 THP-1 cells were transferred to each of the six-well plates and cultured with 100ng/mL phorbol 12-myristate 13-acetate (PMA) (P8139, Sigma-Aldrich) for 24h. Four to five scratches were made in cells within each well.

Human HaCaT cell lines were obtained from the cell library of the Chinese Academy of Sciences and cultured in RPMI medium supplemented with 10% FBS, 100 U/mL penicillin, and 100 mg/mL streptomycin at 37 °C in a 5% CO2 incubator. The influence of AREG was studied by adding different concentrations of recombinant AREG protein (100ng/mL, 500ng/mL, 1000ng/mL) (P15514, novoprotein) to the culture medium for 24h.

### Extraction and copy number determination of cell-free mtDNA

To detect the copy numbers of mtDNA, the DNeasy Blood & Tissue Kit (QIAGEN) was utilized following the manufacturer’s instructions to isolate the cell-free mtDNA in plasma and cell supernatant. Quantitative real-time PCR analyses were conducted to determine the mtDNA copy number in samples, using plasmids expressing mouse or human Cytochrome (CytB) with known copy numbers as standards. The mouse or human CytB sequences were inserted into the pUC57 plasmid, transformed and amplified in Escherichia coli (*E.coli*). The recombinant plasmid DNA was isolated from the bacteria and sequenced to confirm the sequences and acquire the DNA mass. The copy numbers of plasmids were determined by the formula: (6.02×10^23^) (OD26050)/(base number6601014)=copies/μL. Standard plasmids with concentrations of 100ng/μL, 10ng/μL, 1ng/μL, 10-1 ng/μL, 10-2 ng/μL, 10-3 ng/μL, 10-4 ng/μL, 10-5 ng/μL, 10-6 ng/μL were diluted from the plasmids. The samples with standard substances were subjected to RT-PCR simultaneously. The CT-copy number standard curve was calculated using the data of standard plasmids, and the copy numbers of samples were calculated using the same formula.

### Extraction and application of mtDNA

The DNeasy Blood & Tissue Kit (QIAGEN) was used according to the manufacturer’s instructions to extract mtDNA from mouse liver for injection into wound beds. The concentration and purity of mtDNA were determined using the NanoDrop2000c and stored at −20°C. For injection into wound beds, 20μg (mtDNA^lo^_LW) and 80μg (mtDNA^hi^_LW) of mtDNA were used at PWD 0, 3, and 6. For in vitro tests, 100ng/mL mtDNA was added to the cell culture medium for 24h.

### Bulk-RNA sequencing

Bulk-tissue RNA sequencing was conducted using three to four replicates of mice skin wounds in each group using standard methods to make sure samples were strictly controlled for quality. The Illumina sequencing of the libraries was performed. Through z-transformation of fragments per kilobase of transcript per million mapped reads of the selected gene, gene expression was analyzed. Differential expression analysis of two groups (three biological replicates per group) was performed using the DESeq2 R package (v1.32.0). A p value < 0.05 and |log2(foldchange)| > 1.5 were set as thresholds for significant differential expression. We used the cluster Profiler R package to test the statistical enrichment of marker genes in KEGG and GO pathways.

### Single cell RNA sequencing

#### Tissue dissociation

For scRNA-seq analysis, we collected five fresh samples at 2 days post scab detachment (SD2) per group. The wound tissues underwent enzymatic epidermal-dermal separation using the Epidermis Dissociation Kit (Epidermis Dissociation Kit, mouse; Miltenyi Biotec) followed by dissociation of the epidermis and dermis parts. The resulting cells were then filtered, centrifuged, and resuspended in phosphate-buffered saline (PBS) with 0.5% bovine serum albumin (BSA). The dermis part was further dissociated using a mixed enzyme solution containing type I collagenase and trypsin, digested, and then processed similarly to the epidermis part.

Subsequently, the dermis cells were combined with the epidermis cells after being subjected to red blood cell lysis buffer and removal of dead cells and debris by Dead Cell Removal MicroBeads (Miltenyi).

#### Sequencing and data processing

Single-cell suspensions were used for Single-Cell RNA-seq (10x Genomics Chromium Single Cell Kit), followed by sequencing using an Illumina 1.9 mode. Subsequently, the reads were aligned, and expression matrices were generated using the Cell Ranger pipeline software. Downstream computational analysis involved merging different samples into one Seurat object using the RunHarmony function and performing filtering, normalization, scaling, canonical correlation analysis, principal component analysis, and dimensionality reduction using various R packages. Unsupervised clustering and differential gene expression analysis were carried out, followed by gene set enrichment analysis and receptor-ligand probability prediction among cell subpopulations.

#### Pseudotime Analysis

We utilized Monocle 2 for pseudo-temporal trajectory analysis to elucidate cell differentiation trajectories. These sophisticated algorithms position cells along a trajectory that corresponds to a specific biological process, such as cell differentiation, leveraging the distinct asynchronous progression of individual cells within an unsupervised framework(*54, 55*). In the analysis, raw count data of highly variable genes, identified using the FindVariableGenes function from the Seurat package (with parameter y.cutoff = 0.5), were employed for pseudo-temporal trajectory analysis.

#### RNA Velocity Analysis

ScVelo package was employed to perform RNA velocity analysis, which allowed for the identification of transient cellular states and prediction of directional progression of transcriptomic signatures along developmental trajectories. This analysis was based on gene-specific rates of transcription, splicing, and degradation of mRNA, with the results projected as a stream of arrows on the UMAP Embedding.

### Gene signature scoring based on scRNA-seq data

In the assessment of module scores and enrichment fractions for EGF pathway genes and IL17 pathway genes’ expression in individual cells, the scRNA-seq data underwent AddModuleScore() computation. The hallmark genes, associated with the EGF or IL-17 pathway in GO terms, were obtained from the Molecular Signatures Database (http://www.gsea-msigdb.org/gsea/msigdb/index.jsp) and utilized as the gene signature for scoring.

### Histopathology, Immunohistochemistry, and immunofluorescence microscopy

For histopathology, immunohistochemistry, and immunofluorescence microscopy, the samples were fixed with 4% paraformaldehyde and underwent ethanol and xylene dehydration at least 24 hours beforehand. H&E staining and Masson’s trichrome staining were performed to observe re-epithelialization and collagen fiber deposition. In addition, frozen sections were collected for antibodies that can only be stained for IF-Fr. The primary antibodies used for immunofluorescence staining were as follows: TLR9 (Abcam, Ab134368, 1:150), IRF7 (sc-74472, Santa Cruz Biotechnology, 1:133), EEA1 (C45B10, Cell signaling technology, 1:133), CD3 (14-0032-82, Thermo Fisher Scientific, 1:100), Ki67 (Servicebio, GB121141, 1:100), TCR γ/δ (118101, Biolegend, 1:150), Goat Anti-Armenian hamster IgG H&L (Alexa Fluor® 647) (ab173004), CCL2 (26161-1-AP, Proteintech, 1:150), and AREG (16036-1-AP, Proteintech, 1:200). Biopsy sections were de-paraffinized and underwent antigen retrieval using Target Retrieval Solution. After washing and permeabilization with Tris-buffered saline (Quality Biological, 351-086-101) and 0.1% Tween 20 (Sigma, P2287) (TBST) buffer, the sections were blocked with 5% sheep serum and 1% Bovine Serum Albumin (BSA) (Fisher Bioreagents, BP9703-100) at room temperature for 1 hour. Subsequently, the sections were incubated with the primary antibodies (as mentioned above) diluted in Antibody Diluent (Agilent Dako, S0809) overnight at 4°C. After washing with TBST, the sections were incubated with fluorescent binding secondary antibodies at room temperature for 1 hour. Following a final wash, the cell nuclei were stained with DAPI, and the sections were mounted with mounting medium. Immunofluorescent images were analyzed using DFC365FX (Leica) or Olympus FV3000 Confocal Laser Scanning Microscope and processed with FIJI/ImageJ (National Institutes of Health, Bethesda, MD).

### Western Blot Analysis

To perform Western blot analysis, protein lysates were extracted from the wound bed of three different types of snap-frozen skin tissue: WT_LW, TLR9agonist_LW, and enhanced_LW. The proteins (15 µg per lane) were then separated using sodium dodecyl sulfate (SDS)-polyacrylamide gels and transferred to polyvinylidene fluoride membranes. After being blocked with 5% BSA, the membranes were incubated overnight with primary antibodies obtained from Cell Signaling Technology, Inc. (Danvers, MA), which included CCL2 (26161-1-AP, Proteintech, 1:500), AREG (16036-1-AP, Proteintech,1:400), IL-17A(sc-374218, Santa Cruz, 1:500), and GAPDH antibody (sc47724, Santa Cruz, 1:5000). The membrane was then washed and incubated with secondary horse radish peroxidase–labeled antibody. Bands were visualized using FluorChem E (ProteinSimple, San Jose, CA). Densitometry graphs were obtained by quantifying the phosphoprotein and total protein bands through densitometry with the use of Image-Pro Plus software (Media Cybernetics, Inc., Rockville, MD).

#### ALP staining

Whole-mount HFN assay to detect ALP+ dermal papilla (DP) was performed following the protocols in(*1*).

#### Flow cytometry analysis

For flow cytometry analysis, single cells were digested from skin wounds and pre-incubated with purified anti-CD16/CD32 antibody (101301, BioLegend) (1.0 μg per 106 cells in 100 μl volume) for 5 to 10 min to block Fc receptors. The cell suspensions were then co-incubated with fixable viability dye (eFluor™ 780, 65-0865-14, eBioscience) and antibodies against surface markers CD45(PE/Cy7, 147703, BioLegend), Cd3e(APC, 100311, BioLegend), CD4(FITC, 11-0041-82, eBioscience), CD8a(Alexa Fluor™ 700,56-0081-82, eBioscience), and TCRγ/δ(eflour-450,48-5711-82, eBioscience) at 1:400 dilution for 30 min at 4°C in the dark (100 μl per antibody per sample). After fixation and permeabilization, cells were incubated with antibodies against intracellular marker IL-17A (PE,12-7177-81, eBioscience) at 1:400 dilution for 30 min at 4°C in the dark (100 μl per antibody per sample). Fluorescence Minus One (FMO) groups were applied in every test. Flow cytometry analysis was performed using Attune Nxt flow cytometer (Thermo Fisher Scientific) and FlowJo (v10.8.1). The experiments were performed independently three times (n = 3).

#### Statistics and reproducibility

Statistical analyses were performed with Case Viewer software, Image J software, and Prism 9.0 software using two-tailed t-test/ t’-test or one-way analysis of variance (ANOVA) with Tukey post-hoc test. Data were presented as mean ± standard deviation, and a p-value less than 0.05 was considered statistically significant (*p < 0.05, **p < 0.01, ***p < 0.001, ****p < 0.0001), while ns indicates no statistically significant difference.

## Acknowledgments

None.

## Author contributions

Conceptualization: XHL, CH, YM Methodology: XHL, ZMY,SDC, TTA Investigation: XHL, ZMY,SDC, TTA, ZYX, FZ, CD Visualization: XHL Supervision: CH, YM Writing—original draft: XHL Writing—review & editing: XHL, CH

**Competing interests**

All authors declare that they have no competing interests.

**Data and materials availability**

All data to support the conclusions in this manuscript can be found in the figures and the supplementary materials. Other data that support the findings of this study are available from the corresponding author upon reasonable request. After being accepted, RNA sequencing data used in this study are available in the NCBI Sequence Read Archive (SRA) database under accession code.

## Supplementary Materials

**Fig S1.**
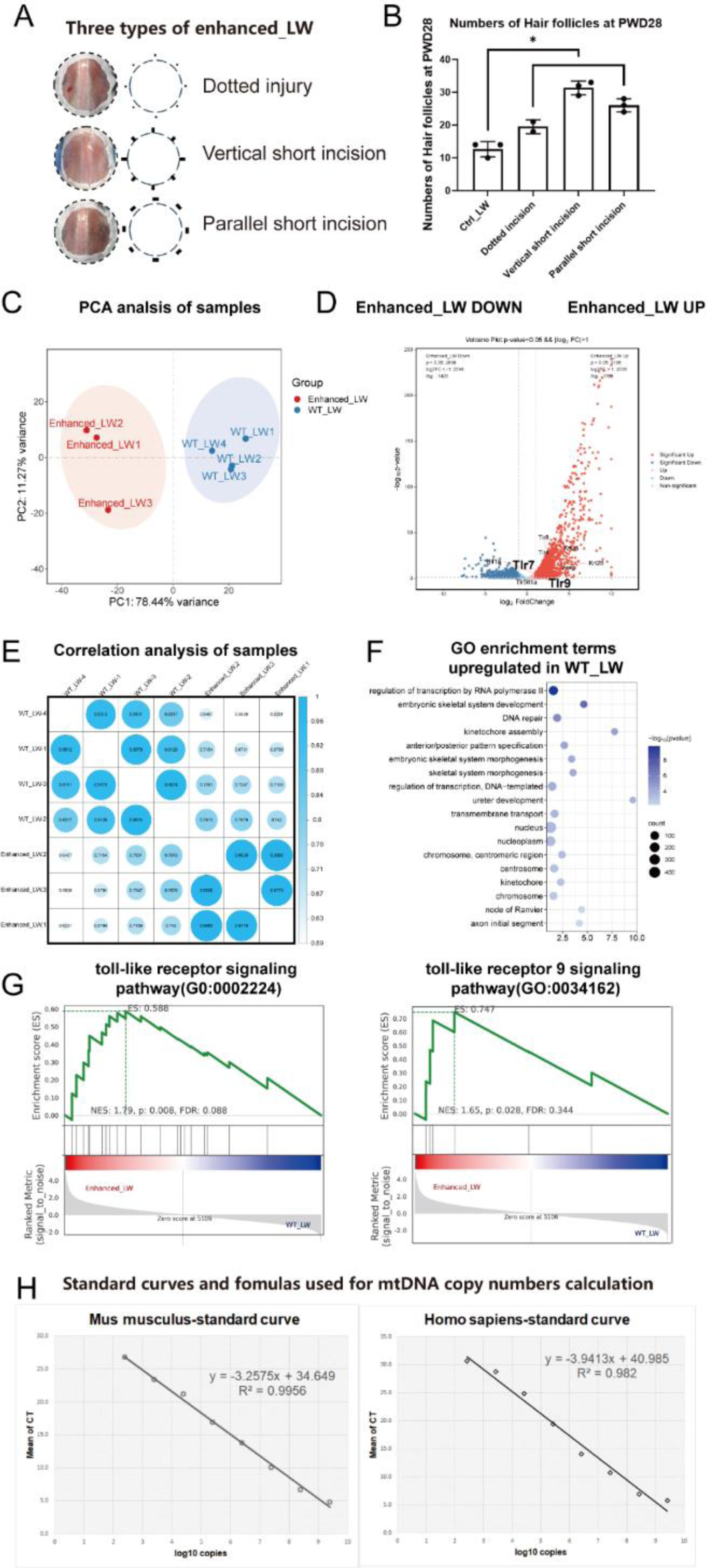
A) Evaluation of the wound healing process with enhanced_LW. A) The three types of enhanced_LW and the wound models. B) Number of regenerated hair follicles in PWD28. Statistical analysis was performed using one-way ANOVA with Dunnett’s multiple comparisons test. n=3 for each group. C) The PCA analysis of bulk-RNA sequence samples. D) The volcano plot showing the genes expressed in enhanced_LW and WT_LW. The threshold for screening differential genes was p-value<0.05 and |log2FC|>1. E) Correlation analysis heatmaps of enhanced_LW and WT_LW. F) GO terms enriched in WT_LW. G) The GSEA enrichment analysis of toll-like receptor signaling pathway and toll-like receptor 9 signaling pathway. H) Standard curves and formulas between CT number and copy numbers calculated by plasmids standards. All error bars ±SD. *p < 0.05, **p < 0.01, ***p < 0.001, and ****p < 0.0001.

**Fig S2.**
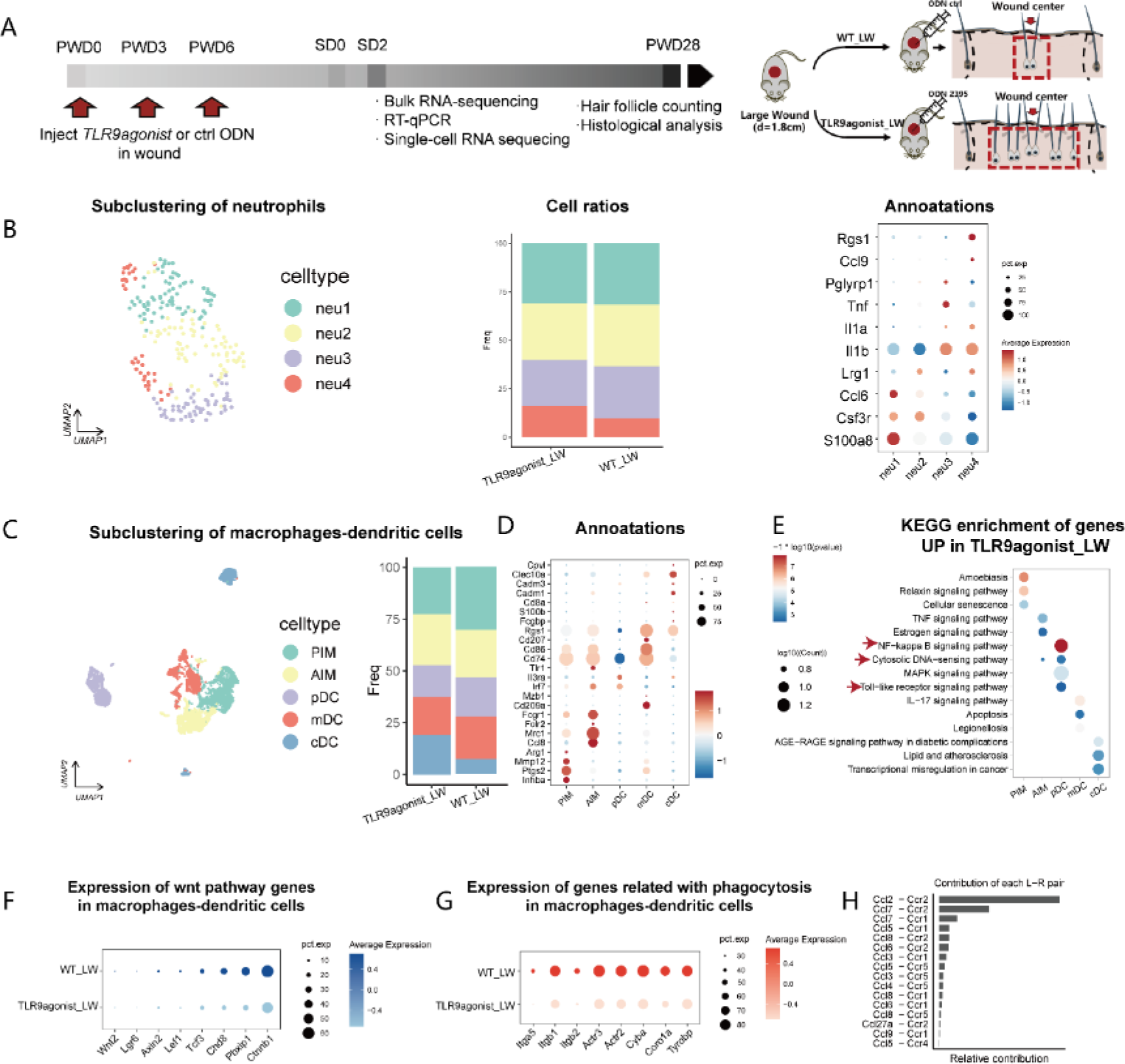
The subcluster and analysis of neutrophils and macrophages. A) Workflow for evaluating the function of TLR9 and mtDNA in large-scale wound healing. The time around scab detachment (SD) was chosen as the key time of hair follicle neogenesis. B) The subclustering of neutrophils and relative cell ratios and marker genes. C) The subclustering of macrophages and dendritic cells. D) The annotations and marker genes of macrophages and dendritic cells. E) The KEGG enrichment terms for every subclusters. F) The expression of wnt pathway genes in macrophages and dendritic cells between two groups. G) The expression of phagocytosis related genes in macrophages and dendritic cells between two groups. H)The contribution rank of each L-R pair in CCL pathway.

**Fig S3.**
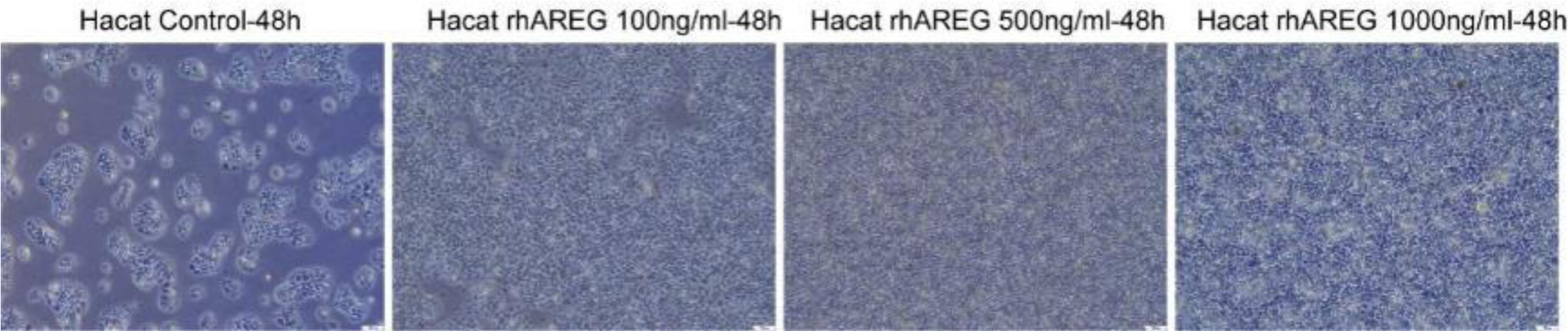
The effect of rhAREG to Hacat after 48h.

**Table S1:**
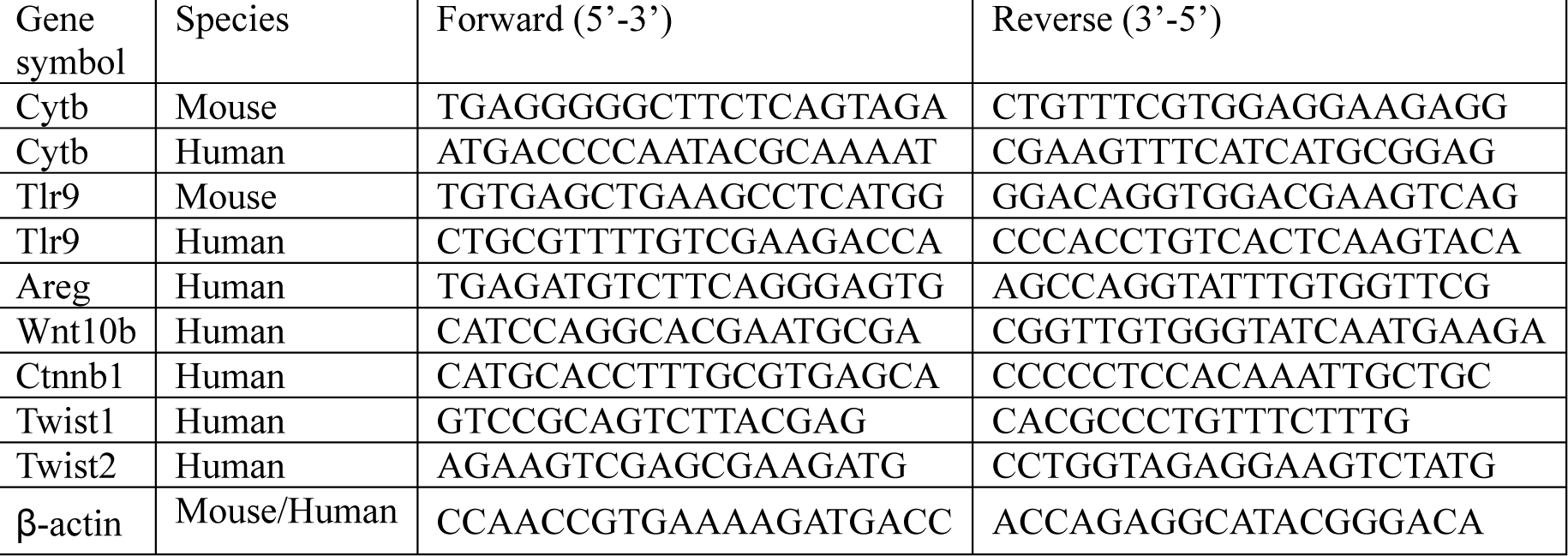
Primers used for RT-qPCR.

